# Screening effects of HCN channel blockers on sleep/wake behavior in zebrafish

**DOI:** 10.1101/2023.05.05.539631

**Authors:** Fusun Doldur-Balli, Sandra Smieszek, Brendan T. Keenan, Amber J. Zimmerman, Olivia J. Veatch, Christos M. Polymeropoulos, Gunther Birznieks, Mihael H. Polymeropoulos

## Abstract

Hyperpolarization-activated cyclic nucleotide-gated (HCN) ion channels generate electrical rhythmicity in various tissues although primarily heart, retina and brain. The HCN channel blocker compound, Ivabradine (Corlanor), is approved by the US Food and Drug Administration (FDA) as a medication to lower heart rate by blocking hyperpolarization activated inward current in the sinoatrial node. In addition, a growing body of evidence suggests a role for HCN channels in regulation of sleep/wake behavior. Zebrafish larvae are ideal model organisms for high throughput drug screening, drug repurposing and behavioral phenotyping studies. We leveraged this model system to investigate effects of three HCN channel blockers (Ivabradine, Zatebradine Hydrochloride and ZD7288) at multiple doses on sleep/wake behavior in wild type zebrafish. Results of interest included shorter latency to day time sleep at 0.1 μM dose of Ivabradine (ANOVA, p:0.02), moderate reductions in average activity at 30 μM dose of Zatebradine Hydrochloride (ANOVA, p:0.024) in daytime, and increased nighttime sleep at 4.5 μM dose of ZD7288 (ANOVA, p:0.036). These differences support the hypothesis that compounds blocking HCN channels decreases wakefulness.

**Highlights:**

- A drug screening study in which effects of HCN channel blocker compounds were tested displayed decreased wakefulwness in zebrafish.
- There was modest evidence of these drugs on sleep and wake phenotypes including shorter latency to sleep, moderate reductions in average activity and increased sleep at different doses of three compounds compared to DMSO.
- While several specific doses of Ivabradine, Zatebradine hydrochloride or ZD7288 demonstrated some differences compared to DMSO, effects of these compounds was smaller than the effect of melatonin, a positive control.

## 1. Introduction

Hyperpolarization-activated cyclic nucleotide-gated (HCN) ion channels are members of the family of the voltage gated ion channels (Sartiani et al. 2017). HCN channels are encoded by the *HCN1-4* gene family (Chang et al. 2019) and can form homotetramers or heterotetramers with specific biophysical properties (Sartiani et al. 2017). These integral membrane proteins (Flynn and Zagotta 2018) generate an inward current in heart (I_f_) and nerve cells (I_h_) (Novella Romanelli et al. 2016). HCN channels are known as pacemakers (Wobig et al. 2020); they modulate cardiac rhythmicity and neuronal excitability (Wobig et al. 2020). Functions of HCN channels in photoreceptors include adaptation of the vertebrate retina to visual stimuli (Barrow and Wu 2009). Notably, HCN channels are also involved in regulation of sleep/wake behavior (Lewis and Chetkovich 2011, Sartiani et al. 2017, Byczkowicz et al. 2019, Chang et al. 2019) by contributing to the formation of spindle waves (McCormick and Pape 1990, Bal and McCormick 1996) and slow wave oscillations during non-Rapid Eye Movement (REM) sleep (Kanyshkova et al. 2009, Zobeiri et al. 2018). There are different reports on how HCN channels fulfill sleep related functions. One line of research suggests that inhibition of HCN channels, thereby inhibition of I_h_ current, via local infusion of melatonin in mouse lateral hypothalamus is associated with reductions in wakefulness (Huang et al. 2020). In contrast, inhibition of I_h_ current via orexin A application to mouse prelimbic cortex increased wakefulness (Li et al. 2010). Another study reported sleep fragmentation in a *Drosophila* mutant model, which lacks I_h_ current; however, no significant difference in total sleep amount was noted between mutant and control flies (Gonzalo-Gomez et al. 2012). These different findings reported in the literature led us to test effects of HCN channel blocker compounds on rest/wake behavior in zebrafish as they are a diurnal vertebrate system for performing high-throughput screening of small molecule compounds. We evaluated Ivabradine (Corlanor), Zatebradine hydrochloride and ZD7288 in this study. Specifically, Ivabradine has been observed to inhibit inward current in cell lines originated from human embryonic kidney cells and Chinese hamster ovary cells and rabbit sinoatrial nodes (Novella Romanelli et al. 2016). Zatebradine inhibited inward current in human embryonic kidney cell lines and Xenopus oocytes (Novella Romanelli et al. 2016). Administration of ZD7288 was found to inhibit inward current in human embryonic kidney cell lines, Chinese hamster ovary cell lines, Xenopus oocytes, rat dorsal ganglion neurons, spontaneously hypertensive ventricular myocytes and Guinea pig sinoatrial nodes (Novella Romanelli et al. 2016). These compounds block HCN subunits nonselectively (Novella Romanelli et al. 2016, Zhong and Darmani 2021). All three compounds are pharmacological tools used to reduce heart rate (Novella Romanelli et al. 2016); however, Ivabradine is the only FDA approved drug used in patients with heart failure (Novella Romanelli et al. 2016). Drug screening studies using zebrafish models have been instrumental in detecting effects of small molecule compounds on regulation of sleep/wake behavior and circadian rhythm (Rihel et al. 2010, Mosser et al. 2019). In addition, zebrafish can be utilized to identify mechanism of action of drugs (Rihel et al. 2010, Hoffman et al. 2016, Mosser et al. 2019). The zebrafish model has several additional advantages, such as yielding a high number of offspring per breeding and high throughput assessment of sleep/wake (Oikonomou and Prober 2017). Sleep phases and regulation of sleep in zebrafish are conserved and meet all the behavioral criteria that are used to define a sleep state (Zhdanova 2006, Rihel et al. 2010). Given these advantages, to reveal effects of HCN channel blocker compounds on sleep/wake behavior, we tested if wild type zebrafish larvae exposed to three compounds, administered at different dosages, expressed differences in multiple sleep-related traits when compared to vehicle (DMSO) exposed fish.

## 2. Experimental Procedures

### 2.1. Zebrafish Sleep/Wake Assay

Larval zebrafish were raised on a 14 h light and 10 h dark cycle at 28.5°C. The entrainment and activity measurement equipment (ViewPoint Life Sciences Inc., aka Zebraboxes) houses 96 well plates and utilizes infrared lights to collect. White light is used to maintain day and night cycle. Recirculating water was utilized to ensure that zebrafish larvae were kept at optimum growth temperature (28.5°C) in the chamber of the equipment. Zebrafish larvae collected from a wild type line (AB line) were individually pipetted into each well of a 96 well plate (Whatman, catalog no: 7701-1651) containing 650 μl of standard E3 embryo medium (5 mM NaCl, 0.17 mM KCl, 0.33 mM CaCl_2_, 0.33 mM MgSO_4_, pH 7.4) at 4 days post fertilization. Embryo medium in the wells was topped off each morning once lights were on. Zebrafish experiments were performed in accordance with University of Pennsylvania Institutional Animal Care and Use Committee (IACUC) guidelines.

### 2.2. Drug Testing

Experiments were performed on 96 well plates. Four wells chosen at random (maximum one per row) did not include any larvae but were instead filled with standard embryo medium (E3 embryo medium) to serve as quality control (QC) for the settings, recording and sensitivity of the equipment. Ivabradine (Cayman, Cas Registry No. 148849-67-6), Zatebradine hydrochloride (Tocris, Cas Registry No. 91940-87-3) and ZD7288 (Tocris, Cas Registry No. 133059-99-1) were tested in this study. Each compound was tested at six concentrations varying between 0.1-30 μM (.i.e., 0.1 μM, 0.3 μM, 1.0 μM, 4.5 μM, 10 μM and 30 μM), as reported previously (Rihel et al. 2010). Each drug was dissolved in DMSO. Stock solutions of Ivabradine, Zatebradine hydrochloride and ZD7288 were prepared at 35 millimolar, 40 millimolar and 30 millimolar concentrations, respectively. As indicated by the manufacturers; solubility of Ivabradine and Zatebradine hydrochloride in DMSO is 20 mg/ml and that of ZD7288 is 100 millimolar. Lower concentrations were obtained by serial dilution. Each dose was tested on 11-12 larvae per plate, depending on the location of the randomly chosen QC wells, for evaluating the impact of different doses of the target drug on sleep and behavioral phenotypes. Zebrafish larvae were allowed to acclimate to the environment by spending the first night without any exposure to drugs and baseline sleep was observed during the second night. Drugs were then added at six days post fertilization at 5:00 pm; this was a one-time drug administration. 96 well plate was removed from the video monitoring equipment to administer drug compound and software continued to capture activity data. The peak in the sleep graph at the time of drug administration was formed when 96 well plate was removed from the equipment. Each assay was performed over a total of four days: acclimation on day 1, tracking baseline sleep on day 2, drug administration on day 3 and data acquisition on days 2-4 (see Figure 1). Concurrently, we studied 11-12 embryos that served as a DMSO (or drug vehicle, 1:1000 vol:vol) exposed control group and 11-12 embryos were exposed to 100 nM (0.1 micromolar) of melatonin as a positive control. Prior literature has utilized this concentration of melatonin to demonstrate sleep-promoting effects (Zhdanova et al. 2001), and our own proof-of-concept data shows that this concentration of melatonin is very effective for increasing sleep in zebrafish larvae (see Supplementary Figure S1). The studies for Ivabradine were repeated six times (three replicates in two Zebraboxes) for a total of 66-72 fish for each drug concentration (11-12 fish per replicate) to ensure robust statistical power. Based on statistical power analysis, providing an effect size of 0.8, appropriate sample size to determine significance was n=25. Therefore, we concluded that three repeats of Zatebradine hydrochloride and ZD7288 assays using a different group of wild type embryos for each replicate would be sufficient by providing three biological replicates for a total of 33-36 fish for each drug concentration (11-12 fish per replicate, all replicates were carried out in the same Zebrabox for each drug).

**Figure 1.**
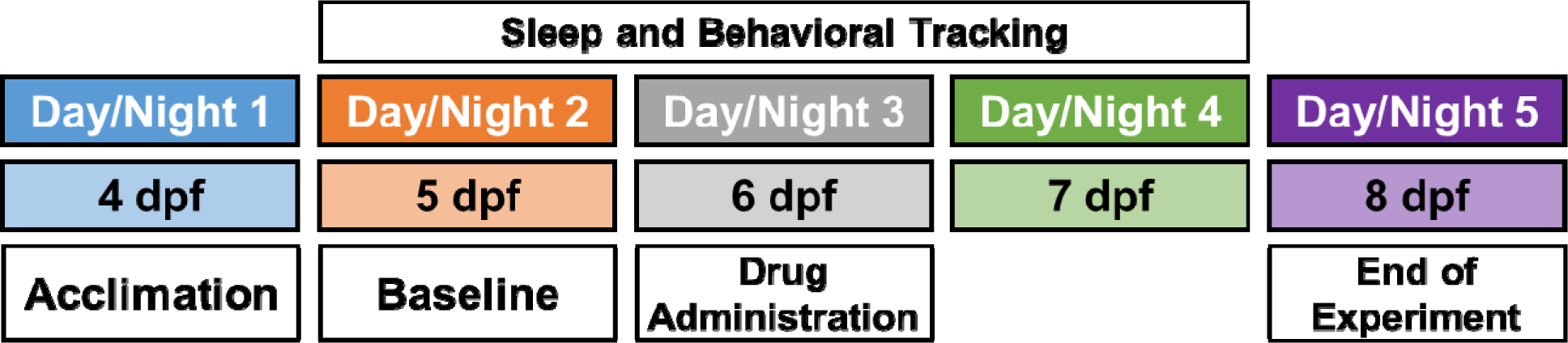
Schematic overview of experimental paradigm, including acclimation on experimental day 1 (4 days post fertilization [dpf]), baseline recording on day 2 (5 dpf), drug administration on day 3 (6 dpf) and sleep and behavioral tracking on days 2-4 (5-7 dpf).

### 2.3. Behavioral Phenotyping

Locomotor activity was captured via a commercially available video tracking system in quantization mode (ViewPoint Life Sciences Inc.) and data were analyzed using a custom designed MATLAB code as previously described (Doldur-Balli et al. 2023). Behavioral tracking took place for three days starting from baseline sleep on day 5 of larval development (see Figure 1). The evaluated sleep phenotypes (see Table 1) included total sleep duration, average activity, average waking activity, sleep bout numbers, consolidation of sleep (average sleep bout length) and latency to sleep as a measure relevant to insomnia. Primary analyses were based on phenotypes calculated within the time window between 30 minutes after drug administration (drug administration was performed at 5:00 pm) and 11 pm (beginning of lights off period). Secondary analyses were performed for the night after drug administration (lights off period).

**Table 1.** Definitions of Key Behavioral Phenotypes in Drug Screening Study

| **Phenotypes** | **Definition** |
| --- | --- |
| **Total Sleep** | Total sleep in minutes. Sleep is defined as any one-minute period of inactivity with less than 0.5 second of total movement of zebrafish larva. |
| **Sleep Bouts** | A continuous sequence of sleep minutes. Total sleep bouts is the bout numbers in a given period. |
| **Sleep Latency** | Length of time (in minutes) spent until the first sleep bout starts. |
| **Bout Length** | Average duration of a sleep bout (in minutes). |
| **Average Activity** | Average activity (seconds/minute) calculated by dividing total activity by total recording minutes |
| **Average Waking Activity** | Average waking activity (seconds/minute) calculated by dividing total activity by the total waking minutes |

### 2.4. Statistical Analysis

Analyses were performed to evaluate phenotypic effects of compounds of interest at six concentrations – 0.1 μM, 0.3 μM, 1.0 μM, 4.5 μM, 10 μM and 30 μM – as reported by Rihel *et al* (Rihel et al. 2010) using complementary approaches. First, to evaluate the relationship between drug doses and sleep phenotypes with minimal assumptions, we performed an analysis of variance (ANOVA) testing whether there were any differences in phenotypes among the experimental groups (DMSO and drug doses). If results for this overall ANOVA were significant (p<0.05), we examined pairwise differences between drug doses and camera-matched DMSO controls to assess which groups were driving the overall differences, including calculation of standardized mean differences (SMDs). The standardized mean difference (SMD) was calculated by dividing the observed mean difference between groups by the pooled standard deviation. As defined by Cohen (Cohen 1988), SMDs of 0.2, 0.5 and 0.8 represent small, medium and large differences, respectively. In addition to ANOVA, two complementary dose-response analyses were performed to evaluate whether a consistent change in sleep phenotypes was observed for increasing drug doses. First, we performed a linear trend analysis, including dose as an ordinal variable in the regression model (e.g., DMSO = 0, 0.1 μM = 1, 0.3 μM = 2, …, 30 μM = 6). This model treats differences between doses as similar in magnitude, asking whether there is a linear increase for higher dosage groups. Second, dose was included as a continuous variable in linear regression, to estimate the expected change in outcome for a 1 μM increase in drug dose; these analyses give increased weight to differences between DMSO and higher dosage groups (e.g. 10 μM or 30 μM). A p-value <0.05 was considered evidence of a significant association across all analyses. To maximize statistical power, analyses were performed pooling data from all experiments. To help account for potential batch effects, the experimental replicate (1, 2 or 3) was included as a covariate and analyses of Ivabradine also included a covariate for experimental box (1 or 2), as two different boxes were utilized. In addition, all analyses performed on data measured after drug administration were adjusted for baseline values of the given phenotype during the same time period prior (i.e., data from the day before and data from the night before were used as baseline values in primary and secondary analysis, respectively). Analyses in which significant associations in both ANOVA and dose-response analyses are observed were considered the most robust evidence for an effect of the drug compound. Results in which there were observed differences based on ANOVA but not following dose-response analyses were assumed to suggest a single dose of drug may be driving the overall results.

### 2.5. Power and Sample Size

Our study included between 33-36 larvae per drug concentration across three biological replicates. This represents nearly twice the maximum sample size utilized by a previous zebrafish drug screening study which detected significant effects (Rihel et al. 2010). Furthermore, for pairwise contrasts, ≥33 animals per group were estimated to provide >80% power to detect standardized effect size differences (i.e., Cohen’s d) of at least 0.70 at an α=0.05, which represent moderate-large effects. Analyses leveraging all data to examine the linear dose response (n ≈ 240 total larvae) were well-powered to detect considerably smaller effects, including >90% power for a correlation of 0.21 (equal to 4.4% variance in sleep behavior explained by drug concentration [R^2^ = 0.044]).

## 3. Results

### 3.1. Summary

Visual inspection of plots of sleep/wake phenotypes across Ivabradine, Zatebradine hydrochloride and ZD7288 doses in some experiments suggested characteristics consistent with increased sleep on the day of drug administration. However, any differences observed with these drug compounds were smaller than the effect of melatonin. Melatonin increased sleep immediately after drug administration (see Figures 2, 3, and 4). Each drug dose was tested on 33-36 zebrafish larvae in three biological replicates. Results of analyses performed as described in Section 2.4 for each drug are presented in more detail below.

**Figure 2.**
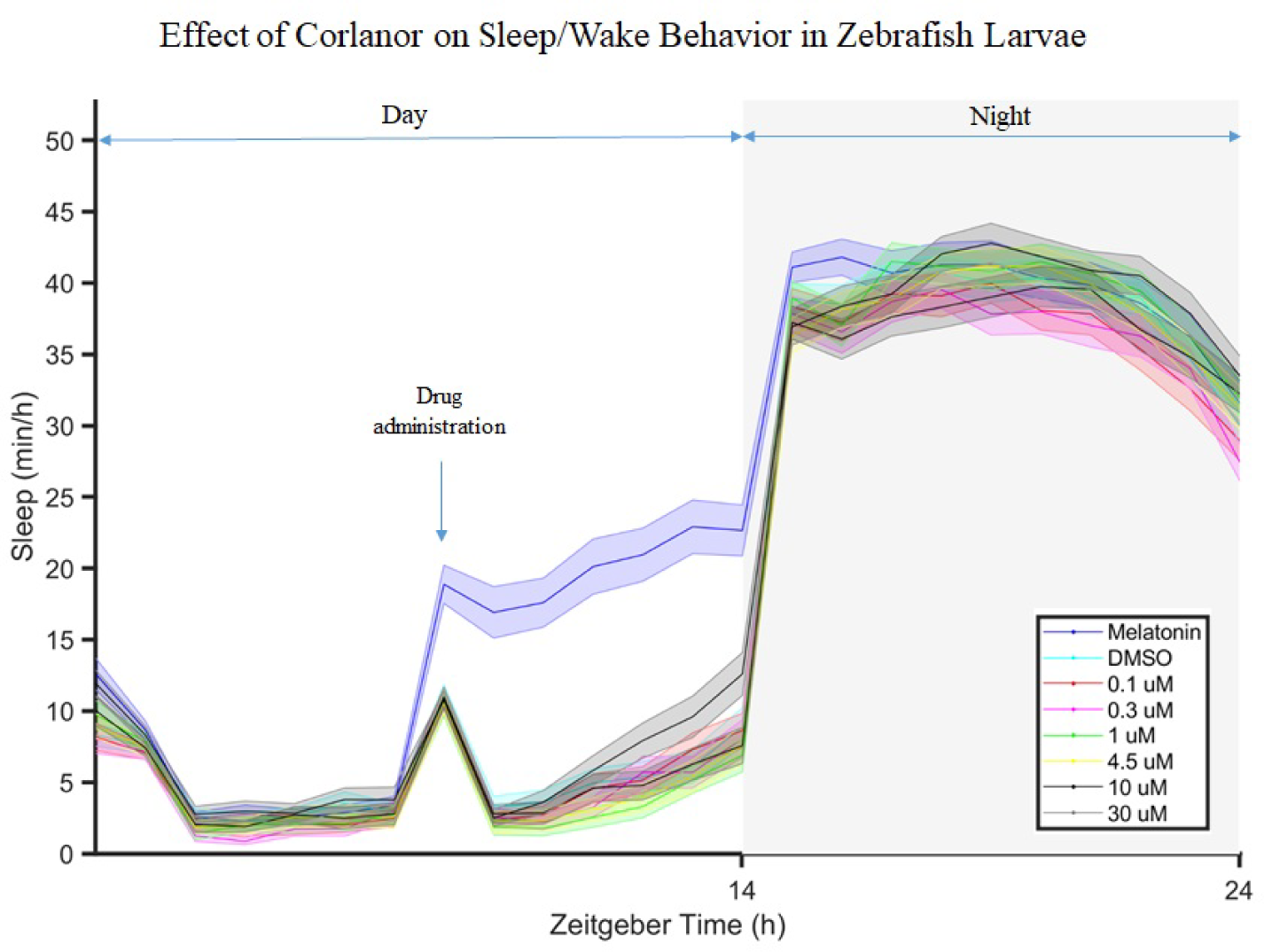
Effect of Ivabradine (Corlanor) on sleep of zebrafish larvae. Different final concentrations of this drug compound ranging between 0.1 μM and 30 μM, DMSO control and 0.1 μM melatonin were added to wells containing 6 days post fertilization larvae. Trending but non-significant increase in sleep duration, likely driven by the higher mean value in the 30 μM Ivabradine group can be observed. Total n:550, melatonin n:67, DMSO n:68, 0.1 μM n:69, 0.3 μM n:71, 1 μM n:66, 4.5 μM n:69, 10 μM n:69, 30 μM n:71, n:number of animals.

**Figure 3.**
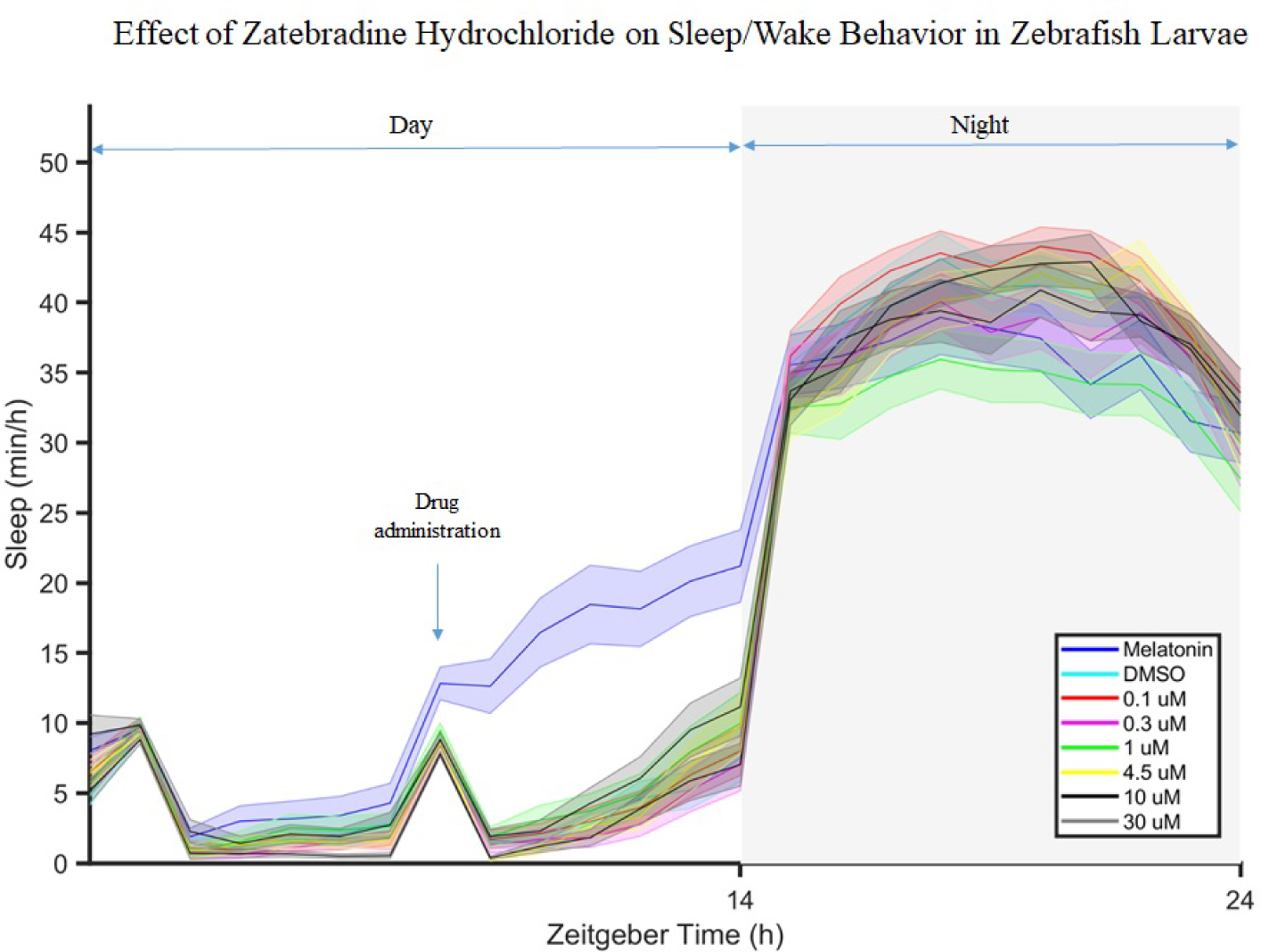
Effect of Zatebradine hydrochloride on sleep of zebrafish larvae. Different final concentrations of this drug compound ranging between 0.1 μM and 30 μM, DMSO control and 0.1 μM melatonin were added to wells containing 6 days post fertilization larvae. The 30 μM dose group showed significantly lower activity compared to DMSO. Total n:275, melatonin n:35, DMSO n:34, 0.1 μM n:34, 0.3 μM n:34, 1 μM n:35, 4.5 μM n:35, 10 μM n:34, 30 μM n:34, n:number of animals.

**Figure 4.**
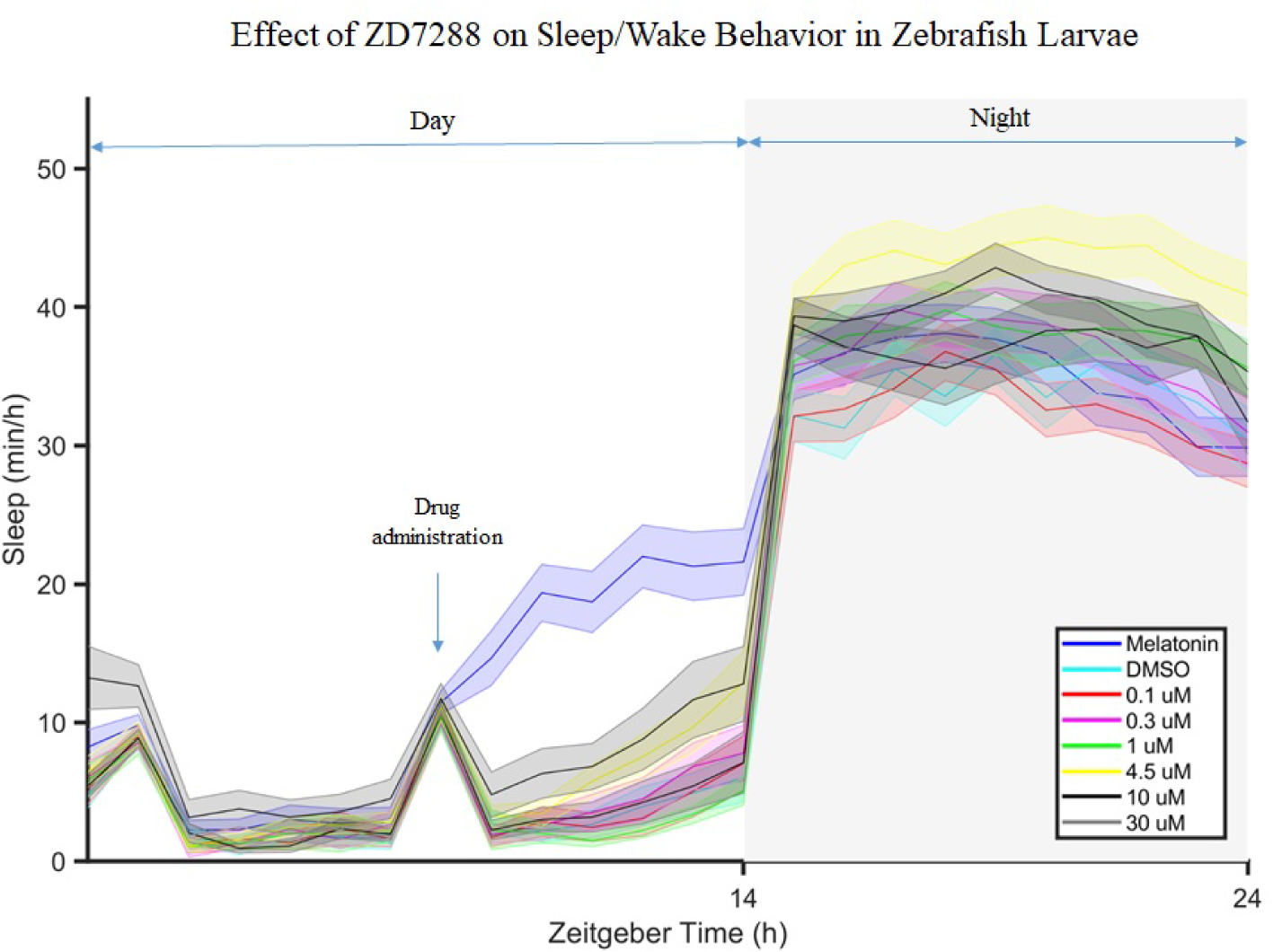
Effect of ZD7288 on sleep of zebrafish larvae. Different final concentrations 3 of this drug compound ranging between 0.1 μM and 30 μM, DMSO control and 0.1 μM 4 melatonin were added to wells containing 6 days post fertilization larvae. There is some 5 visual evidence that ZD7288 increased sleep at 4.5 and 30 μM doses after drug was 6 administered but overall ANOVA was not significant. Total sleep was significantly 7 increased at 4.5 μM dose compared to DMSO during lights off period. Total n:275, 8 melatonin n:35, DMSO n:35, 0.1 μM n:34, 0.3 μM n:35, 1 μM n:34, 4.5 μM n:34, 10 9 μM n:34, 30 μM n:34, n:number of animals.

### 3.2. Statistical Analysis of Ivabradine Screening

#### 3.2.1. Primary Analysis

Primary analyses of phenotypes as calculated within the time window between 30 minutes after Ivabradine administration (drug administration was performed at 5:00 pm) and 11 pm (beginning of the lights off period) are presented in Table 2. In ANOVA comparisons among groups, there was a difference in sleep latency (p = 0.020), with a shorter latency in the 0.1 μM group compared to DMSO (SMD = -0.321, p = 0.048). No differences in latency were observed between DMSO and other dosage groups, and results of linear and continuous dosage models were non-significant (see Table 2). Near significant differences—following ANOVA—were observed in average activity (p = 0.094), and average waking activity (p = 0.073). For both endpoints, continuous dosage models suggested some decreased activity for each 1 μM increase in Ivabradine, likely driven by the lower mean value in the 30 μM group. For comparison to differences between DMSO and Ivabradine doses, results of analyses comparing DMSO to the positive control melatonin during the same time period are shown in Supplementary Table S1. Strong differences between DMSO and melatonin were observed for all phenotypes (all p≤0.006), with absolute standardized mean differences (SMDs) ranging from 0.49 for bout length to 1.15 for average waking activity.

**Table 2.** Sleep and Activity Immediately after Drug Administration across Ivabradine Doses

| Phenotype | Adjusted Mean (95% CI)<sup>†</sup> |  |  |  |  |  |  | ANOVA<br>p<sup>‡</sup> | Linear Model<sup>§</sup> |  | Dosage Model<sup>¶</sup> |  |
| --- | --- | --- | --- | --- | --- | --- | --- | --- | --- | --- | --- | --- |
| | DMSO | 0.1 μM | 0.3 μM | 1.0 μM | 4.5 μM | 10 μM | 30 μM | | $\beta$ (95% CI) | p | $\beta$ (95% CI) | p |
| **Total Sleep, minutes** | 30.2<br>(21.8, 38.6) | 33.7<br>(25.3, 42.1) | 29.6<br>(21.3, 37.8) | 22.2<br>(13.7, 30.8) | 26.0<br>(17.6, 34.4) | 30.2<br>(21.9, 38.6) | 34.3<br>(26.0, 42.6) | 0.447 | 0.08<br>(-1.50, 1.66) | 0.924 | 0.18<br>(-0.13, 0.49) | 0.253 |
| **Sleep Bouts, number** | 13.1<br>(10.2, 16.1) | 14.2<br>(11.2, 17.1) | 12.8<br>(9.9, 15.7) | 11.6<br>(8.6, 14.6) | 10.3<br>(7.3, 13.2) | 13.3<br>(10.4, 16.2) | 15.8<br>(12.9, 18.7) | 0.217 | 0.13<br>(-0.43, 0.69) | 0.649 | 0.10<br>(-0.01, 0.21) | 0.073 |
| **Sleep Latency, minutes** | 127.8<br>(101.2, 154.4) | **89.9**<br>**(63.6, 116.3)\*** | 150.8<br>(124.8, 176.9) | 148.6<br>(121.6, 175.6) | 114.8<br>(88.4, 141.2) | 123.6<br>(97.2, 149.9) | 111.4<br>(85.3, 137.5) | **0.020** | -0.67<br>(-5.70, 4.36) | 0.793 | -0.53<br>(-1.52, 0.458) | 0.293 |
| **Bout Length, minutes** | 2.03<br>(1.74, 2.31) | 2.03<br>(1.76, 2.30) | 2.20<br>(1.91, 2.49) | 1.83<br>(1.52, 2.14) | 1.99<br>(1.71, 2.27) | 1.93<br>(1.64, 2.21) | 2.02<br>(1.75, 2.30) | 0.775 | -0.015<br>(-0.068, 0.038) | 0.578 | -0.001<br>(-0.011, 0.010) | 0.901 |
| **Avg. Activity, sec/min** | 3.26<br>(3.05, 3.47) | 3.27<br>(3.06, 3.48) | 3.40<br>(3.19, 3.60) | 3.33<br>(3.12, 3.54) | 3.37<br>(3.16, 3.58) | 3.22<br>(3.01, 3.43) | 2.96<br>(2.75, 3.17) | 0.094 | -0.036<br>(-0.077, 0.006) | 0.094 | **-0.012**<br>**(-0.020, -0.004)** | **0.003** |
| **Avg. Wake Act., sec/min** | 3.32<br>(3.12, 3.53) | 3.37<br>(3.16, 3.58) | 3.48<br>(3.28, 3.68) | 3.38<br>(3.17, 3.59) | 3.45<br>(3.25, 3.65) | 3.29<br>(3.08, 3.49) | 3.04<br>(2.83, 3.24) | 0.073 | -0.037<br>(-0.077, 0.004) | 0.075 | **-0.012**<br>**(-0.020, -0.004)** | **0.002** |
| Statistically significant associations (p<0.05) shown in **bold**; <sup>†</sup>Model estimated mean and 95% confidence interval, adjusted for replicate, experimental box, and baseline values of phenotype; <sup>‡</sup>p-value from ANOVA testing whether there are any differences in phenotype among dosage groups; <sup>§</sup>Results from linear model treating dose as an ordinal variable – $\beta$ represents the expected change in phenotype associated with increasing to the next highest dosage group; <sup>¶</sup>Results from continuous dosage model – $\beta$ represents the expected change in phenotype for 1 μM increase in dose; *p<0.05 compared to DMSO in pairwise comparisons (performed only when ANOVA p<0.05). | | | | | | | | | | | | |

#### 3.2.2. Secondary Analysis

Secondary analysis was performed for sleep phenotypes during the lights off period after Ivabradine administration (Supplementary Table S2). No significant differences among Ivabradine doses were observed based on ANOVA. A small increase in total sleep was observed in the continuous dosage model, with an increase of 0.71 minutes (95% CI: 0.09, 1.34) sleep for each 1 μM increase in Ivabradine (p = 0.025). Results comparing DMSO and melatonin are again presented as a positive control (Supplementary Table S3). Small to moderate differences were observed with Melatonin in the number (SMD = -0.49, p = 0.002) and length (SMD = 0.38, p = 0.007) of sleep bouts, but there were no differences in total sleep or sleep latency.

### 3.3. Statistical Analysis of Zatebradine Hydrochloride Screening

#### 3.3.1. Primary Analysis

Comparisons of sleep and activity patterns among drug doses immediately after Zatebradine Hydochloride administration are presented in Table 3. Differences were observed among groups for average activity (p = 0.024) and average waking activity (p = 0.030), but there were no differences in other phenotypes based on ANOVA. Compared to DMSO, the 30 μM dose group showed significantly lower average activity (SMD = -0.43, p = 0.032) and average waking (SMD = -0.40, p = 0.041) activity. These differences are reflected in significant associations in continuous dose models for each phenotype, but only trending results in linear models (Table 3). An association (p = 0.034) in the dosage model was also observed for sleep latency, with each 1 μM increase in Zatebradine Hydochloride associated with a 1.61 minute decrease (95% CI: -3.09, -0.13). We again observed significant differences in all phenotypes when comparing DMSO to melatonin as a positive control (Supplementary Table S4), with absolute SMDs ranging from 0.53 for sleep bout length to 1.42 for average activity and average waking activity.

**Table 3.**
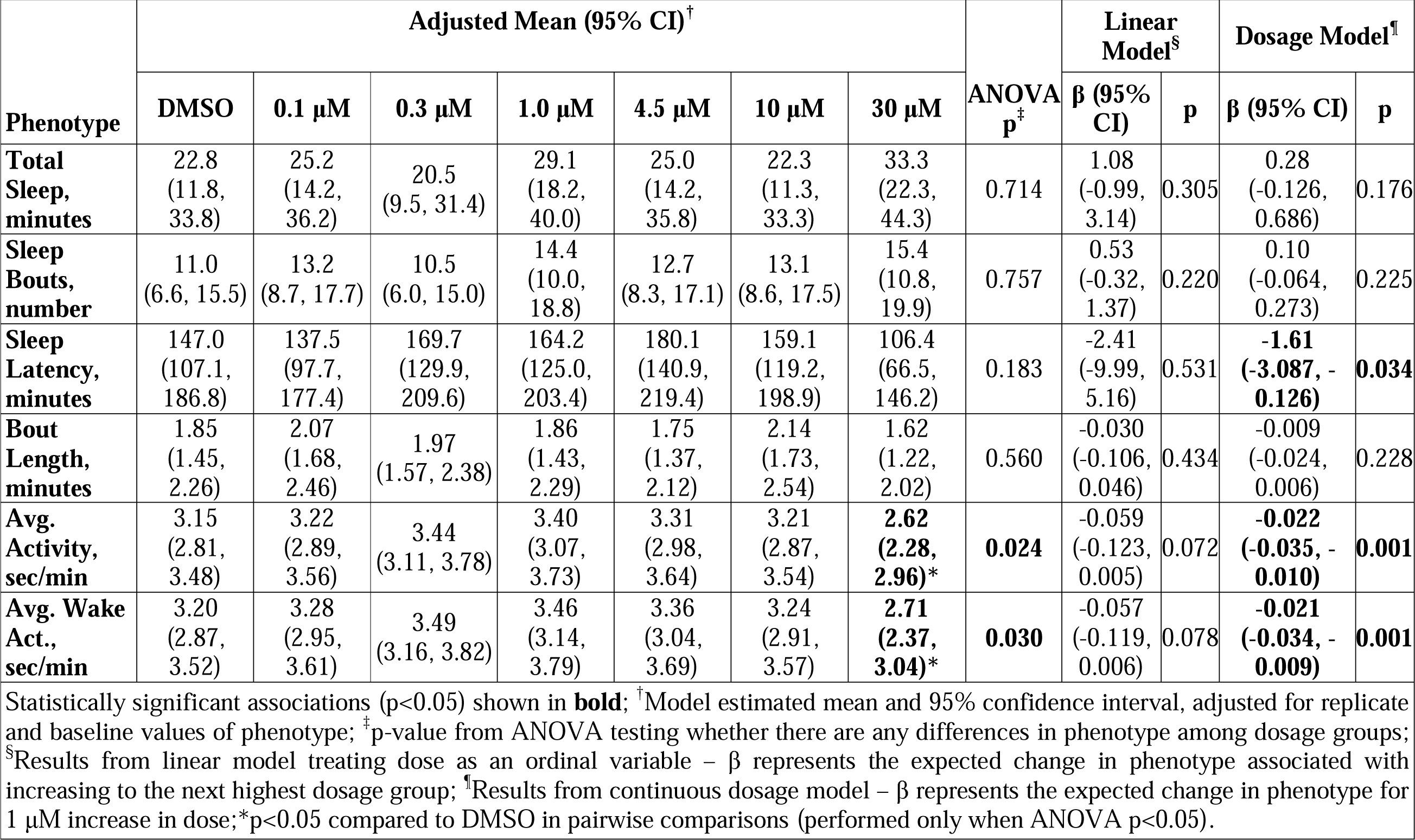
Sleep and Activity Immediately after Drug Administration across Zatebradine Hydrochloride Doses

| Phenotype | Adjusted Mean (95% CI)<sup>†</sup> |  |  |  |  |  |  | ANOVA<br>p<sup>‡</sup> | Linear Model<sup>§</sup> |  | Dosage Model<sup>¶</sup> |  |
| --- | --- | --- | --- | --- | --- | --- | --- | --- | --- | --- | --- | --- |
|  | DMSO | 0.1 µM | 0.3 µM | 1.0 µM | 4.5 µM | 10 µM | 30 µM |  | β (95% CI) | p | β (95% CI) | p |
| **Total Sleep, minutes** | 22.8<br>(11.8, 33.8) | 25.2<br>(14.2, 36.2) | 20.5<br>(9.5, 31.4) | 29.1<br>(18.2, 40.0) | 25.0<br>(14.2, 35.8) | 22.3<br>(11.3, 33.3) | 33.3<br>(22.3, 44.3) | 0.714 | 1.08<br>(-0.99, 3.14) | 0.305 | 0.28<br>(-0.126, 0.686) | 0.176 |
| **Sleep Bouts, number** | 11.0<br>(6.6, 15.5) | 13.2<br>(8.7, 17.7) | 10.5<br>(6.0, 15.0) | 14.4<br>(10.0, 18.8) | 12.7<br>(8.3, 17.1) | 13.1<br>(8.6, 17.5) | 15.4<br>(10.8, 19.9) | 0.757 | 0.53<br>(-0.32, 1.37) | 0.220 | 0.10<br>(-0.064, 0.273) | 0.225 |
| **Sleep Latency, minutes** | 147.0<br>(107.1, 186.8) | 137.5<br>(97.7, 177.4) | 169.7<br>(129.9, 209.6) | 164.2<br>(125.0, 203.4) | 180.1<br>(140.9, 219.4) | 159.1<br>(119.2, 198.9) | 106.4<br>(66.5, 146.2) | 0.183 | -2.41<br>(-9.99, 5.16) | 0.531 | **-1.61**<br>**(-3.087, -0.126)** | **0.034** |
| **Bout Length, minutes** | 1.85<br>(1.45, 2.26) | 2.07<br>(1.68, 2.46) | 1.97<br>(1.57, 2.38) | 1.86<br>(1.43, 2.29) | 1.75<br>(1.37, 2.12) | 2.14<br>(1.73, 2.54) | 1.62<br>(1.22, 2.02) | 0.560 | -0.030<br>(-0.106, 0.046) | 0.434 | -0.009<br>(-0.024, 0.006) | 0.228 |
| **Avg. Activity, sec/min** | 3.15<br>(2.81, 3.48) | 3.22<br>(2.89, 3.56) | 3.44<br>(3.11, 3.78) | 3.40<br>(3.07, 3.73) | 3.31<br>(2.98, 3.64) | 3.21<br>(2.87, 3.54) | **2.62**<br>**(2.28, 2.96)\*** | **0.024** | -0.059<br>(-0.123, 0.005) | 0.072 | **-0.022**<br>**(-0.035, -0.010)** | **0.001** |
| **Avg. Wake Act., sec/min** | 3.20<br>(2.87, 3.52) | 3.28<br>(2.95, 3.61) | 3.49<br>(3.16, 3.82) | 3.46<br>(3.14, 3.79) | 3.36<br>(3.04, 3.69) | 3.24<br>(2.91, 3.57) | **2.71**<br>**(2.37, 3.04)\*** | **0.030** | -0.057<br>(-0.119, 0.006) | 0.078 | **-0.021**<br>**(-0.034, -0.009)** | **0.001** |

Statistically significant associations (p<0.05) shown in **bold**; <sup>†</sup>Model estimated mean and 95% confidence interval, adjusted for replicate and baseline values of phenotype; <sup>‡</sup>p-value from ANOVA testing whether there are any differences in phenotype among dosage groups; <sup>§</sup>Results from linear model treating dose as an ordinal variable – β represents the expected change in phenotype associated with increasing to the next highest dosage group; <sup>¶</sup>Results from continuous dosage model – β represents the expected change in phenotype for 1 µM increase in dose; \*p<0.05 compared to DMSO in pairwise comparisons (performed only when ANOVA p<0.05).

#### 3.3.2. Secondary Analysis

Secondary analyses were performed for sleep phenotypes in the lights off period after Zatebradine Hydochloride administration (Supplementary Table S5). There were no among group differences based on ANOVA. There was statistically significant (p = 0.025) evidence of a small increase in the number of sleep bouts for a 1 μM increase in dosage. There were no differences between DMSO and Melatonin in the lights off period for these phenotypes in this experiment (Supplementary Table S6).

### 3.4. Statistical Analysis of ZD7288 Screening

#### 3.4.1. Primary Analysis

Comparisons of sleep and activity patterns across doses immediately after ZD7288 administration are presented in Table 4. No differences were observed based on ANOVA results. In dose response analyses of bout length, both the linear model and dosage model indicated longer bouts with increased dose of ZD7288 (p = 0.021 and p=0.005). The linear model showed an increased bout length of 0.13 minutes per increase in dosage group (p = 0.021) and the dosage model showed an increased bout length of 0.03 minutes per 1 μM increase (p = 0.005). As in previous experiments, comparisons between DMSO and melatonin as a positive control demonstrated significant differences across all phenotypes (Supplementary Table S7), with absolute SMDs ranging from 0.82 for sleep bout length to 1.52 for the number of sleep bouts.

**Table 4.**
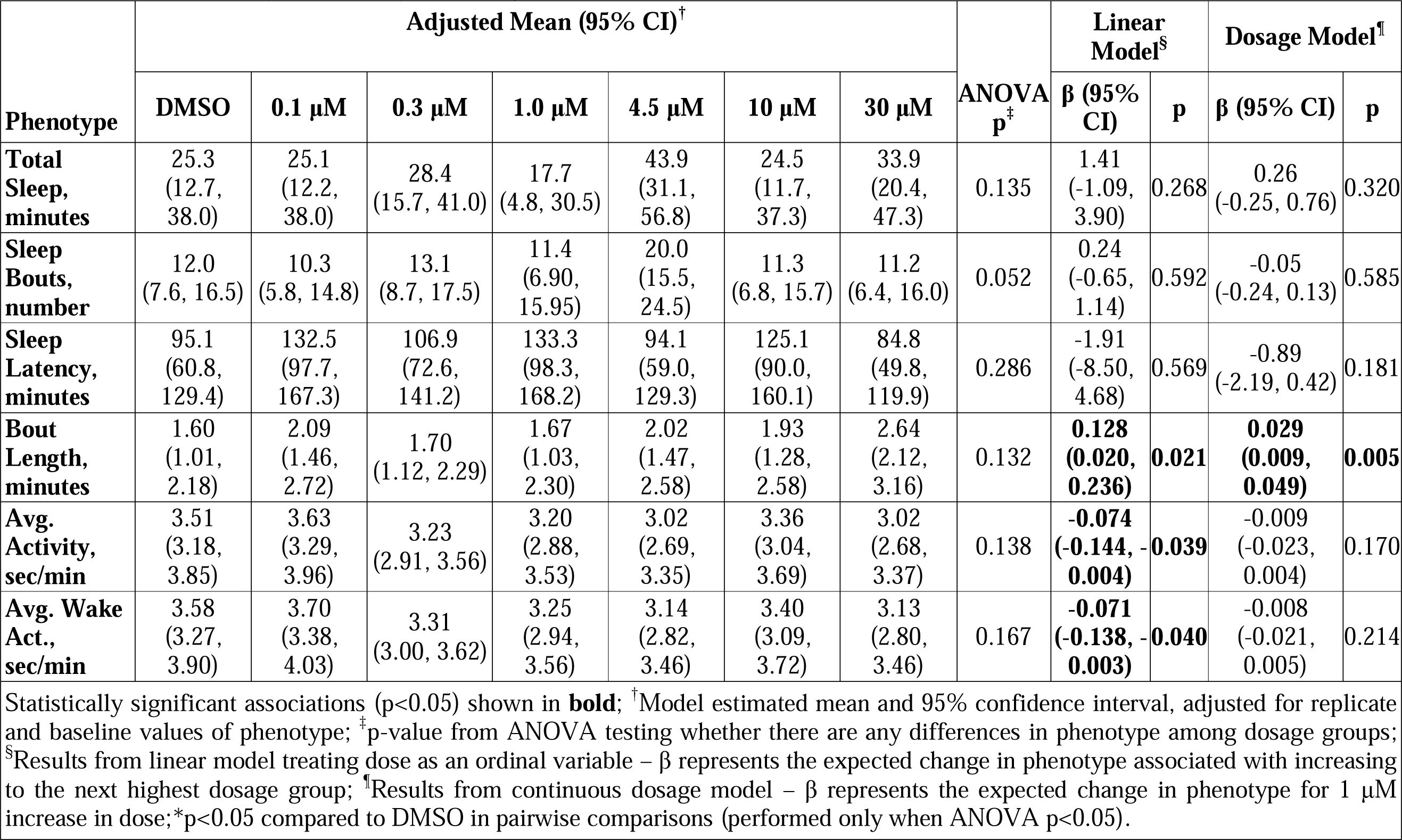
Sleep and Activity Immediately after Drug Administration across ZD7288 Doses

#### 3.4.2. Secondary Analysis

Secondary analyses were performed for sleep phenotypes in the lights off period after ZD7288 administration (Supplementary Table S8). In ANOVA comparisons, differences among dosage groups were observed for total sleep (p = 0.036), number of sleep bouts (p = 0.0003) and sleep bout length (p = 0.003); there was a near significant difference in sleep latency (p = 0.064). Interestingly, differences among groups were driven by an increase in total sleep (SMD = 0.53, p = 0.008), decreased number of sleep bouts (SMD = -0.75, p = 0.001), and increased sleep bout length (SMD = 0.54, p = 0.003) within the 4.5 μM group compared to DMSO. In addition, the 10 μM group demonstrated fewer sleep bouts than DMSO (SMD = -0.49, p = 0.023). These associations between sleep phenotypes and moderate doses of ZD7288 are reflected in significant associations in the linear dose response analyses (Supplementary Table S8). There were no differences between DMSO and melatonin for these phenotypes in the lights off period (Supplementary Table S9).

## 4. Discussion

Screening of HCN channel blockers on sleep/wake behavior of zebrafish larvae showed shorter latency to sleep at 0.1 μM dose of Ivabradine, moderate reductions in average activity at 30 μM dose of Zatebradine Hydrochloride, and increased sleep at 4.5 μM dose of ZD7288 as a result of ANOVA analysis in our study. Among these results, reduction in activity following Zatebradine Hydrochoride administration was supported by dosage model and increased sleep following ZD7288 administration was supported by linear model. All of our findings associated with different sleep parameters were in the same direction. Three tested HCN channel blocker compounds decreased wakefulness. There were different reports on effects of blockade of HCN channels such as decreased (Huang, Li and Leng 2020) and increased wakefulness (Li et al., 2010) in mouse models, fragmented sleep (Gonzalo-Gomez et al., 2012) and no change in total sleep duration (Gonzalo-Gomez et al., 2012) in a *Drosophila* model. The differences we found were in support of the reports suggesting that blocking HCN channels decreases wakefulness.

Waking activity data is utilized to assess health status of the zebrafish larvae in sleep/wake assays (Rihel et al. 2010). We didn’t see large changes in waking activity following administration of HCN channel blockers. This indicates that the doses administered were not toxic and zebrafish larvae were healthy during the assessed period of time.

The half-life of the three HCN channel blocker compounds range between two-three hours (Valenzuela et al. 1996, Chaplan et al. 2003, Tse and Mazzola 2015). Compounds were administered at 5 pm and our primary analysis took place between 05:30-11 pm. Thus our primary analysis included distribution half-life of the tested three HCN channel blockers. Ivabradine does not cross the blood brain barrier (Savelieva and Camm 2008). Zatebradine hydrochloride passes blood brain barrier (Kruger et al. 2000). Ability of ZD7288 to pass blood brain barrier is not known (Zhong and Darmani 2021). Blood brain barrier is sealed by day 5 into development in zebrafish (O’Brown et al. 2019). We administered HCN channel blockers to zebrafish larvae at 6 dpf. Thus, we mimicked the conditions of how humans take HCN channel blockers in our study.

Zatebradine hydrochloride inhibits inward current in Purkinje cells (Valenzuela et al. 1996). Cerebellar activity was lower in NREM sleep compared to that in wakefulness and it was reported to be elevated during REM sleep (Zhang et al. 2020). Also Purkinje cells were active before transitioning from sleep to wakefulness (Zhang et al. 2020). Reduction in activity following Zatebradine hydrochloride administration in the current study points out decreased wakefulness and this finding is in line with the aforementioned reports, most likely through a mechanism affecting Purkinje cells. Moreover, HCN channel blocker compounds including Ivabradine (Demontis et al. 2009), Zatebradine hydrochloride (Satoh and Yamada 2002) and ZD7288 (Satoh and Yamada 2000) inhibit Ih in HCN channels especially HCN1 in rod photoreceptors which contributes to photoreceptor degeneration (Schon et al. 2016). Given that zebrafish are highly responsive to light in sleep regulation (Jones 2007), blockade of photoreceptors in retina might have a role in decreasing wakefulness in our study.

Use of zebrafish as a model organism provided us the opportunity to assess effects of compounds on the whole brain instead of focusing on one brain region at a time (Huang, Li and Leng 2020; Li et al., 2010). Zebrafish is a diurnal organism like humans. This was another advantage of using zebrafish over using a mouse model as mice are nocturnal. *Drosophila* is an invertebrate model (Gonzalo-Gomez et al., 2012). Since zebrafish is a vertebrate model, it possesses more evolutionarily conserved features with mammals compared to *Drosophila* such as nervous system and neuropharmacology (Oikonomou and Prober 2017). Zebrafish is an attractive *in vivo* model to perform drug repurposing studies (Cousin et al. 2014, Wittmann et al. 2015, Sourbron et al. 2019). In this study, we tested effects of HCN channel blocker compounds, which are used as pharmacological tools to reduce heart rate, on sleep/wake behaviors in zebrafish larvae. Our findings which show effects of all three HCN channel blocker compounds in the same direction (ie. decreased wakefulness) can be utilized to determine time of drug intake. In addition, blocking HCN channels has been suggested to be effective in pain treatment (Ramirez et al. 2018). Zebrafish drug screening libraries (Rihel et al. 2010) can be also utilized to identify the pathways through which HCN channel blocker compounds exert their functions associated with alleviating neuropathic pain.

Limitations of our model of choice might be due to the method of drug administration. Drug compounds solved in DMSO were pipetted into individual wells of a 96 well plate in which individual larva swims rather than directly administering it such as injecting. However this is the standard method of drug screening in zebrafish (Rihel et al. 2010, Mosser et al. 2019).

Our study was the first report of testing effects of HCN channel blockers in zebrafish to our knowledge. We also displayed and analyzed effects of melatonin in zebrafish larvae. Ivabradine, Zatebradine hydrochloride and ZD7288 do not work selectively on HCN channel subunits (Novella Romanelli et al. 2016). Each HCN channel subunit might be targeted genetically in future studies to dissect the role of each gene in sleep/wake behavior.

## Supplementary Figures and Tables

**Figure S1.**
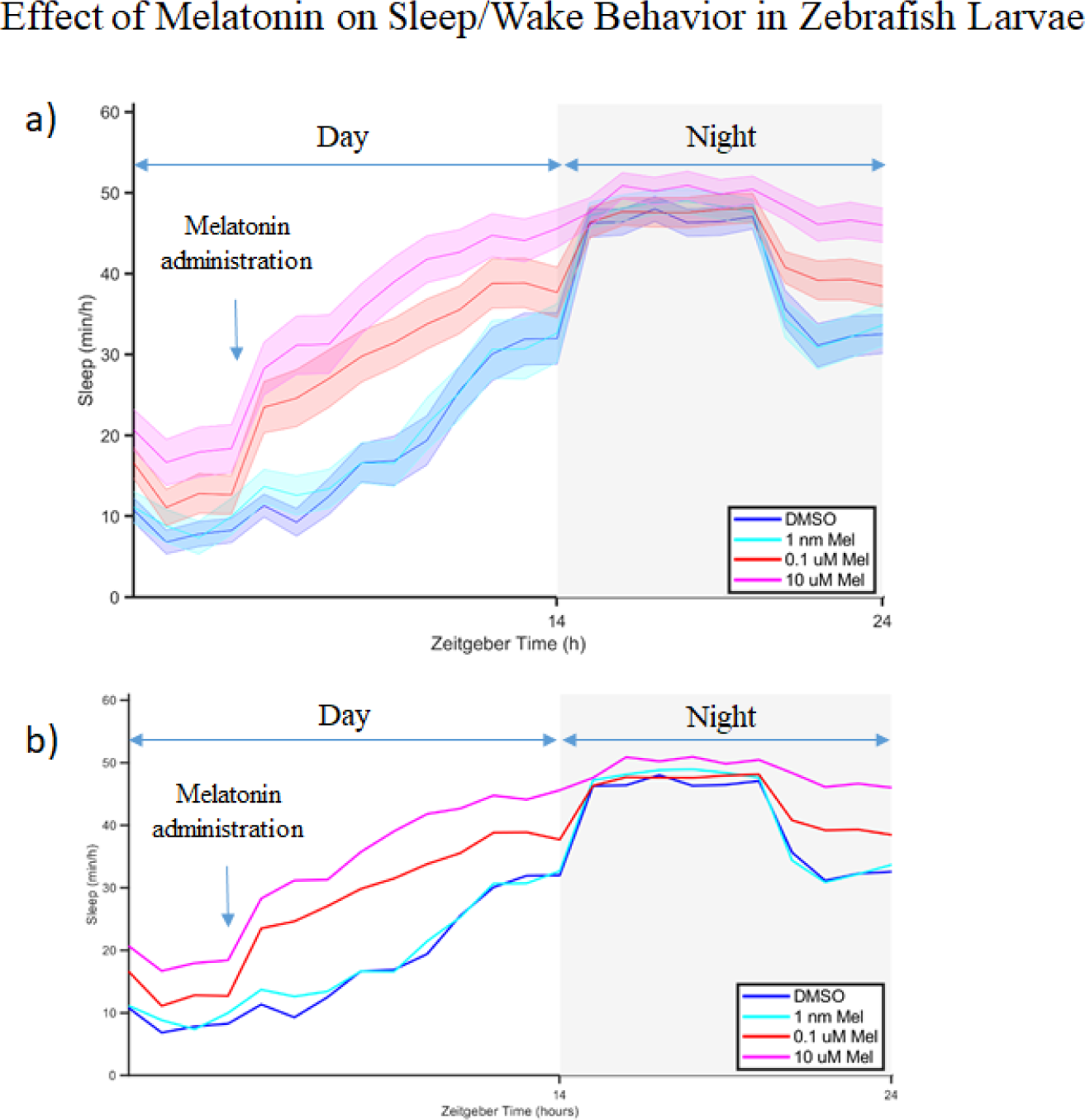
Effect of melatonin on sleep of zebrafish larvae. Three different final concentrations of melatonin (1 nanomolar, 100 nanomolar (0.1 micromolar) and 10 micromolar) and DMSO control were added to the wells containing 6 days post fertilization larvae. a) Sleep graph of 1 hour bins with error bars. b) Sleep graph of 1 hour bins without error bars. There is a robust increase in sleep immediately after melatonin administration at concentrations over 1 nanomolar. n:45 per group.

**Table S1.** Sleep and Activity Immediately After Ivabradine Administration For Melatonin as a Positive Control*

| Phenotype | Adjusted Mean (95% CI)<sup>†</sup> |  | Difference (95% CI)<sup>‡</sup> | p<sup>§</sup> |
| --- | --- | --- | --- | --- |
|  | DMSO | Melatonin |  |  |
| **Total Sleep, minutes** | 35.6 (22.1, 49.1) | 116.5 (102.9, 130.1) | **80.9 (61.7, 100.1)** | **<0.0001** |
| **Sleep Bouts, number** | 14.7 (10.4, 19.0) | 41.4 (37.1, 45.8) | **26.7 (20.5, 32.8)** | **<0.0001** |
| **Sleep Latency, minutes** | 121.0 (99.5, 142.5) | 21.7 (0.0, 43.4) | **-99.3 (-129.9, -68.7)** | **<0.0001** |
| **Bout Length, minutes** | 2.04 (1.69, 2.39) | 2.73 (2.40, 3.06) | **0.69 (0.21, 1.17)** | **0.006** |
| **Avg. Activity, sec/min** | 3.22 (3.01, 3.42) | 1.28 (1.07, 1.48) | **-1.94 (-2.24, -1.64)** | **<0.0001** |
| **Avg. Wake Act., sec/min** | 3.30 (3.10, 3.51) | 1.35 (1.14, 1.56) | **-1.95 (-2.25, -1.65)** | **<0.0001** |
| Statistically significant differences (p<0.05) shown in **bold**; <sup>†</sup>Model estimated mean and 95% confidence interval, adjusted for replicate, experimental box, and baseline values of phenotype; <sup>‡</sup>Adjusted mean difference and 95% CI between melatonin and DMSO groups; <sup>§</sup>p-value comparing phenotype between DMSO and Melatonin groups. *These data are from experiments that involved studies of Ivabradine. |  |  |  |  |

**Table S2.** Sleep during the lights off period after Drug Administration of different doses of Ivabradine and with DMSO as a control

| Phenotype | Adjusted Mean (95% CI)<sup>†</sup> |  |  |  |  |  |  | ANOVA p<sup>‡</sup> | Linear Model<sup>§</sup> |  | Dosage Model<sup>¶</sup> |  |
| --- | --- | --- | --- | --- | --- | --- | --- | --- | --- | --- | --- | --- |
|  | DMSO | 0.1 µM | 0.3 µM | 1.0 µM | 4.5 µM | 10 µM | 30 µM |  | β (95% CI) | p | β (95% CI) | p |
| **Total Sleep, minutes** | 377.5<br>(360.4, 394.6) | 375.2<br>(358.2, 392.2) | 366.6<br>(349.9, 383.3) | 384.4<br>(367.1, 401.8) | 377.5<br>(360.6, 394.5) | 366.5<br>(349.5, 383.5) | 398.9<br>(382.1, 415.6) | 0.118 | 2.12<br>(-1.08, 5.33) | 0.194 | **0.71**<br>**(0.09, 1.34)** | **0.025** |
| **Sleep Bouts, number** | 80.6<br>(76.9, 84.3) | 74.5<br>(70.8, 78.2) | 74.4<br>(70.7, 78.0) | 78.4<br>(74.6, 82.1) | 78.2<br>(74.6, 81.9) | 78.8<br>(75.1, 82.5) | 80.7<br>(77.1, 84.4) | 0.071 | 0.47<br>(-0.23, 1.17) | 0.185 | 0.13<br>(-0.007, 0.267) | 0.063 |
| **Sleep Latency, minutes** | 6.86<br>(6.05, 7.68) | 5.77<br>(4.95, 6.58) | 6.26<br>(5.46, 7.07) | 5.67<br>(4.84, 6.50) | 6.84<br>(6.02, 7.65) | 6.90<br>(6.09, 7.72) | 6.41<br>(5.61, 7.22) | 0.160 | 0.054<br>(-0.100, 0.208) | 0.489 | 0.010<br>(-0.021, 0.040) | 0.532 |
| **Bout Length, minutes** | 5.02<br>(4.56, 5.48) | 5.29<br>(4.84, 5.75) | 5.32<br>(4.87, 5.77) | 5.15<br>(4.69, 5.62) | 5.20<br>(4.75, 5.66) | 5.05<br>(4.60, 5.51) | 5.58<br>(5.13, 6.03) | 0.678 | 0.038<br>(-0.047, 0.124) | 0.380 | 0.012<br>(-0.005, 0.028) | 0.176 |

Statistically significant associations (p<0.05) shown in **bold**; <sup>†</sup>Model estimated mean and 95% confidence interval, adjusted for replicate, experimental box, and baseline values of phenotype; <sup>‡</sup>p-value from ANOVA testing whether there are any differences in phenotype among dosage groups; <sup>§</sup>Results from linear model treating dose as an ordinal variable – β represents the expected change in phenotype associated with increasing to the next highest dosage group; <sup>¶</sup>Results from continuous dosage model – β represents the expected change in phenotype for 1 µM increase in dose; \*p<0.05 compared to DMSO in pairwise comparisons (performed only when ANOVA p<0.05).

**Table S3.** Sleep during the lights off period after Ivabradine administration for Melatonin as a positive control*

| Phenotype | Adjusted Mean (95% CI)<sup>†</sup> |  | Difference (95% CI)<sup>‡</sup> | p<sup>§</sup> |
| --- | --- | --- | --- | --- |
|  | DMSO | Melatonin |  |  |
| **Total Sleep, minutes** | 379.7 (362.9, 396.5) | 394.7 (377.7, 411.6) | 15.0 (-8.9, 38.8) | 0.218 |
| **Sleep Bouts, number** | 81.0 (77.2, 84.8) | 72.5 (68.7, 76.3) | **-8.5 (-13.9, -3.1)** | **0.002** |
| **Sleep Latency, minutes** | 6.75 (5.95, 7.54) | 6.50 (5.69, 7.30) | -0.25 (-1.38, 0.89) | 0.667 |
| **Bout Length, minutes** | 5.01 (4.53, 5.49) | 5.96 (5.47, 6.44) | **0.95 (0.27, 1.63)** | **0.007** |
| Statistically significant differences (p<0.05) shown in **bold**; <sup>†</sup>Model estimated mean and 95% confidence interval, adjusted for replicate, experimental box, and baseline values of phenotype; <sup>‡</sup>Adjusted mean difference and 95% CI between melatonin and DMSO groups; <sup>§</sup>p-value comparing phenotype between DMSO and Melatonin groups. * These data are from experiments that involved studies of Ivabradine. |  |  |  |  |

**Table S4.**
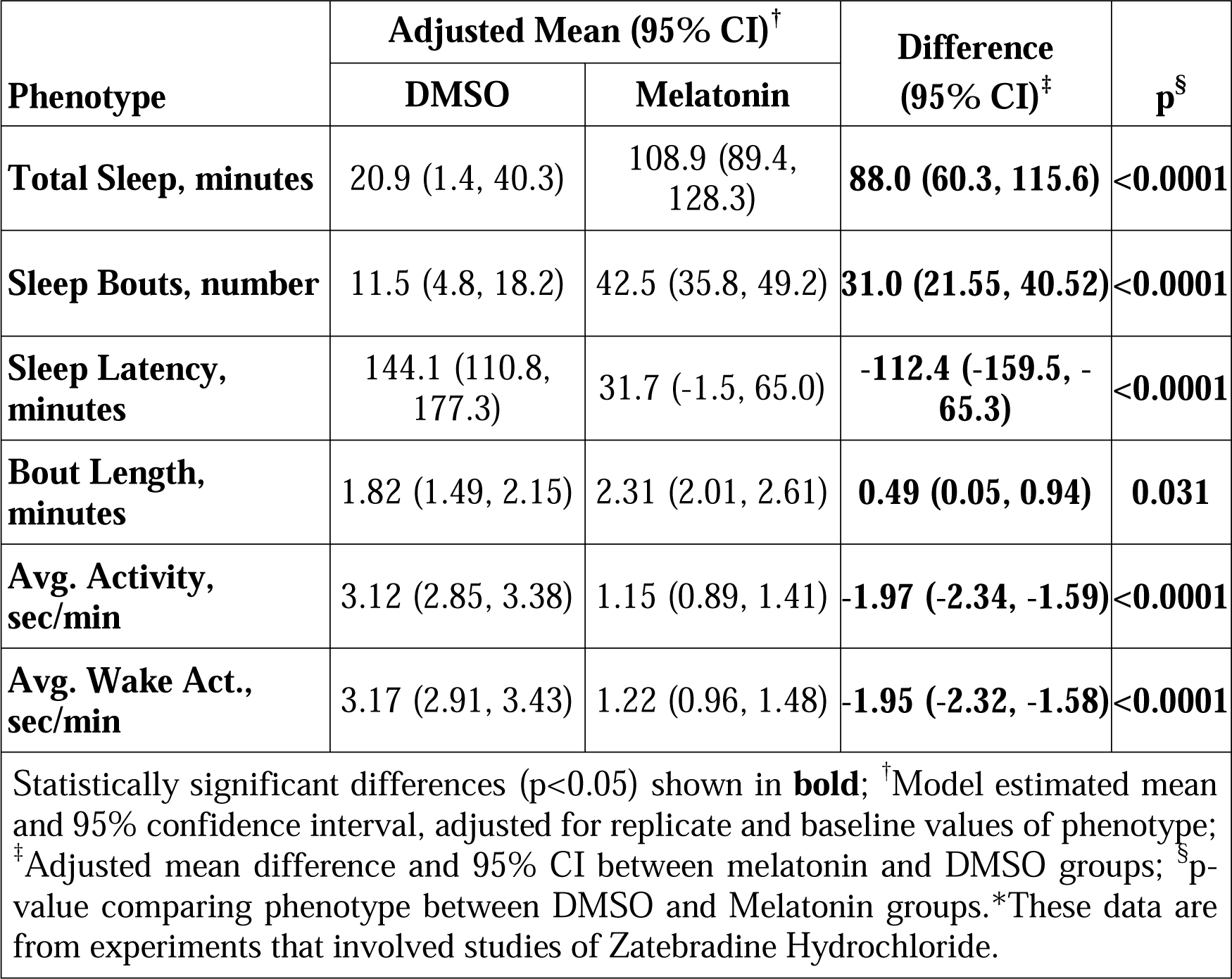
Sleep and Activity Immediately after Zatebradine Hydrochloride Administration for Melatonin As a Positive Control*

**Table S5.** Sleep during the lights off period after Drug Administration of different doses of Zatebradine Hydrochloride and with DSMO as a control

| Phenotype | Adjusted Mean (95% CI)<sup>†</sup> |  |  |  |  |  |  | ANOVA<br>p<sup>‡</sup> | Linear Model<sup>§</sup> |  | Dosage Model<sup>¶</sup> |  |
| --- | --- | --- | --- | --- | --- | --- | --- | --- | --- | --- | --- | --- |
|  | DMSO | 0.1 µM | 0.3 µM | 1.0 µM | 4.5 µM | 10 µM | 30 µM |  | β (95% CI) | p | β (95% CI) | p |
| **Total Sleep, minutes** | 386.2<br>(359.3, 413.2) | 393.7<br>(366.4, 421.0) | 369.4<br>(342.5, 396.4) | 337.1<br>(310.5, 363.7) | 384.6<br>(357.9, 411.2) | 377.5<br>(350.6, 404.4) | 388.6<br>(361.7, 415.5) | 0.068 | -0.25<br>(-5.45, 4.95) | 0.92<br>5 | 0.52<br>(-0.50, 1.53) | 0.31<br>9 |
| **Sleep Bouts, number** | 78.2<br>(73.0, 83.3) | 79.0<br>(73.8, 84.1) | 76.0<br>(70.9, 81.1) | 76.7<br>(71.7, 81.8) | 77.1<br>(72.1, 82.2) | 82.8<br>(77.7, 87.9) | 83.3<br>(78.2, 88.5) | 0.264 | 0.87<br>(-0.10, 1.83) | 0.07<br>9 | **0.22**<br>**(0.03, 0.41)** | **0.02**<br>**5** |
| **Sleep Latency, minutes** | 7.45<br>(5.94, 8.97) | 8.29<br>(6.78, 9.81) | 7.87<br>(6.36, 9.38) | 8.93<br>(7.43, 10.44) | 7.72<br>(6.23, 9.22) | 7.11<br>(5.59, 8.62) | 7.02<br>(5.51, 8.53) | 0.582 | -0.139<br>(-0.424, 0.146) | 0.33<br>8 | -0.039<br>(-0.095, 0.016) | 0.16<br>6 |
| **Bout Length, minutes** | 5.28<br>(4.63, 5.93) | 5.35<br>(4.69, 6.01) | 5.11<br>(4.47, 5.76) | 4.90<br>(4.26, 5.54) | 5.27<br>(4.63, 5.91) | 4.74<br>(4.09, 5.39) | 5.08<br>(4.43, 5.72) | 0.841 | -0.059<br>(-0.181, 0.063) | 0.34<br>1 | -0.005<br>(-0.029, 0.019) | 0.66<br>7 |
| Statistically significant associations (p<0.05) shown in **bold**; <sup>†</sup>Model estimated mean and 95% confidence interval, adjusted for replicate and baseline values of phenotype; <sup>‡</sup>p-value from ANOVA testing whether there are any differences in phenotype among dosage groups; <sup>§</sup>Results from linear model treating dose as an ordinal variable – β represents the expected change in phenotype associated with increasing to the next highest dosage group; <sup>¶</sup>Results from continuous dosage model – β represents the expected change in phenotype for 1 µM increase in dose; *p<0.05 compared to DMSO in pairwise comparisons (performed only when ANOVA p<0.05). |  |  |  |  |  |  |  |  |  |  |  |  |

**Table S6.** Sleep during the lights off period after Zatebradine Hydrochloride Administration for Melatonin As a Positive Control*

| Phenotype | Adjusted Mean (95% CI)<sup>†</sup> |  | Difference<br>(95% CI)<sup>‡</sup> | p<sup>§</sup> |
| --- | --- | --- | --- | --- |
|  | DMSO | Melatonin |  |  |
| **Total Sleep,**<br>**minutes** | 382.6 (353.7,<br>411.5) | 358.0 (329.2,<br>386.9) | -24.6 (-65.6,<br>16.5) | 0.237 |
| **Sleep Bouts,**<br>**number** | 79.2 (73.8, 84.7) | 78.7 (73.2, 84.1) | -0.5 (-8.3, 7.2) | 0.892 |
| **Sleep Latency,**<br>**minutes** | 7.34 (5.44, 9.24) | 8.55 (6.65,<br>10.45) | 1.21 (-1.48,<br>3.90) | 0.372 |
| **Bout Length,**<br>**minutes** | 5.19 (4.65, 5.73) | 4.59 (4.05, 5.13) | -0.60 (-1.37,<br>0.17) | 0.123 |
| <sup>†</sup>Model estimated mean and 95% confidence interval, adjusted for replicate and baseline values of phenotype; <sup>‡</sup>Adjusted mean difference and 95% CI between melatonin and DMSO groups; <sup>§</sup>p-value comparing phenotype between DMSO and Melatonin groups.* These data are from experiments that involved studies of Zatebradine Hydrochloride. |  |  |  |  |

**Table S7.** Sleep and Activity Immediately after ZD7288 Administration for Melatonin As a Positive Control*

| Phenotype | Adjusted Mean (95% CI)<sup>†</sup> |  | Difference<br>(95% CI)<sup>‡</sup> | p<sup>§</sup> |
| --- | --- | --- | --- | --- |
|  | DMSO | Melatonin |  |  |
| **Total Sleep, minutes** | 22.5 (4.6, 40.4) | 110.3 (92.4, 128.2) | **87.8 (62.2, 113.3)** | **<0.0001** |
| **Sleep Bouts, number** | 10.6 (5.7, 15.5) | 44.3 (39.4, 49.2) | **33.7 (26.7, 40.7)** | **<0.0001** |
| **Sleep Latency, minutes** | 94.7 (69.9, 119.5) | 12.4 (-12.4, 37.2) | **-82.3 (-117.7, -46.9)** | **<0.0001** |
| **Bout Length, minutes** | 1.60 (1.24, 1.95) | 2.39 (2.07, 2.71) | **0.80 (0.32, 1.28)** | **0.002** |
| **Avg. Activity, sec/min** | 3.82 (3.50, 4.14) | 1.26 (0.94, 1.57) | **-2.56 (-3.03, -2.10)** | **<0.0001** |
| **Avg. Wake Act., sec/min** | 3.90 (3.59, 4.20) | 1.37 (1.07, 1.68) | **-2.52 (-2.97, -2.08)** | **<0.0001** |
| Statistically significant differences (p<0.05) shown in **bold**; <sup>†</sup>Model estimated mean and 95% confidence interval, adjusted for replicate and baseline values of phenotype; <sup>‡</sup>Adjusted mean difference and 95% CI between melatonin and DMSO groups; <sup>§</sup>p-value comparing phenotype between DMSO and Melatonin groups.<br>*These data are from experiments that involved studies of ZD7288. |  |  |  |  |

**Table S8.** Sleep during the lights off period after Drug Administration with different doses of ZD7288

| Phenotype | Adjusted Mean (95% CI)<sup>†</sup> |  |  |  |  |  |  | ANO<br>VA p<sup>‡</sup> | Linear Model<sup>§</sup> |  | Dosage Model<sup>¶</sup> |  |
| --- | --- | --- | --- | --- | --- | --- | --- | --- | --- | --- | --- | --- |
| | DMSO | 0.1 μM | 0.3 μM | 1.0 μM | 4.5 μM | 10 μM | 30 μM | | $\beta$ (95% CI) | p | $\beta$ (95% CI) | p |
| **Total Sleep, minutes** | 356.8<br>(328.6, 385.0) | 340.1<br>(311.7, 368.5) | 371.1<br>(343.3, 398.8) | 374.8<br>(346.6, 402.9) | **412.1**<br>**(383.4, 440.8)\*** | 365.7<br>(337.6, 393.9) | 383.1<br>(354.8, 411.5) | **0.036** | **5.75**<br>**(0.23, 11.27)** | **0.041** | 0.56<br>(-0.52, 1.63) | 0.310 |
| **Sleep Bouts, number** | 78.0<br>(72.7, 83.2) | 81.1<br>(75.8, 86.4) | 72.4<br>(67.1, 77.6) | 73.1<br>(67.8, 78.4) | **64.6**<br>**(59.2, 69.9)\*** | **69.1**<br>**(63.8, 74.4)\*** | 77.0<br>(71.6, 82.3) | **0.0003** | **-1.24**<br>**(-2.27, -0.21)** | **0.019** | 0.02<br>(-0.19, 0.22) | 0.867 |
| **Sleep Latency, minutes** | 7.12<br>(5.99, 8.25) | 8.11<br>(6.96, 9.26) | 5.63<br>(4.50, 6.76) | 7.12<br>(5.97, 8.27) | 6.05<br>(4.90, 7.20) | 6.44<br>(5.29, 7.59) | 6.38<br>(5.22, 7.53) | 0.064 | -0.181<br>(-0.400, 0.038) | 0.104 | -0.020<br>(-0.063, 0.024) | 0.378 |
| **Bout Length, minutes** | 5.18<br>(4.42, 5.95) | 4.57<br>(3.80, 5.34) | 5.49<br>(4.73, 6.25) | 5.38<br>(4.61, 6.15) | **6.87**<br>**(6.10, 7.64)\*** | 6.13<br>(5.36, 6.90) | 5.53<br>(4.76, 6.29) | **0.003** | **0.195**<br>**(0.047, 0.343)** | **0.010** | 0.012<br>(-0.017, 0.042) | 0.418 |

<sup>†</sup>Model estimated mean and 95% confidence interval, adjusted for replicate and baseline values of phenotype; <sup>‡</sup>p-value from ANOVA testing whether there are any differences in phenotype among dosage groups; <sup>§</sup>Results from linear model treating dose as an ordinal variable – $\beta$ represents the expected change in phenotype associated with increasing to the next highest dosage group; <sup>¶</sup>Results from continuous dosage model – $\beta$ represents the expected change in phenotype for 1 μM increase in dose; \*p<0.05 compared to DMSO in pairwise comparisons (performed only when ANOVA p<0.05).

**Table S9.** Sleep during the lights off period after ZD7288 Administration for Melatonin As a Positive Control*

| Phenotype | Adjusted Mean (95% CI)<sup>†</sup> |  | Difference (95% CI)<sup>‡</sup> | p<sup>§</sup> |
| --- | --- | --- | --- | --- |
|  | DMSO | Melatonin |  |  |
| **Total Sleep, minutes** | 348.3 (318.0, 378.5) | 337.5 (307.2, 367.7) | -10.8 (-54.3, 32.7) | 0.621 |
| **Sleep Bouts, number** | 80.5 (74.6, 86.4) | 78.2 (72.2, 84.1) | -2.3 (-10.7, 6.1) | 0.582 |
| **Sleep Latency, minutes** | 7.05 (5.74, 8.36) | 7.38 (6.07, 8.69) | 0.33 (-1.53, 2.19) | 0.728 |
| **Bout Length, minutes** | 4.56 (3.90, 5.22) | 4.59 (3.93, 5.26) | 0.04 (-0.90, 0.97) | 0.939 |
| <sup>†</sup>Model estimated mean and 95% confidence interval, adjusted for replicate and baseline values of phenotype; <sup>‡</sup>Adjusted mean difference and 95% CI between melatonin and DMSO groups; <sup>§</sup>p-value comparing phenotype between DMSO and Melatonin groups.* These data are from experiments that involved studies of ZD7288. |  |  |  |  |

